# Automated Specimen Triage for Dark Taxa: Deep Learning Enables Orientation, Sex Identification, and Anatomical Segmentation from Robotic Imaging

**DOI:** 10.1101/2025.10.02.680063

**Authors:** Hossein Shirali, Lorenz Wührl, Leshon Lee, Nathalie Klug, Rudolf Meier, Christian Pylatiuk, Emily Hartop

## Abstract

Robotic specimen processing is transforming biodiversity discovery by replacing labor-intensive handling with scalable systems that can simultaneously generate high-quality specimen images. We demonstrate that these images can be leveraged by deep learning to efficiently extract key biological information and guide targeted specimen processing. Using a model dark taxon, the Phoridae (Diptera), the workflow performs three core tasks: sex identification, specimen orientation classification, and anatomical segmentation. Sex identification allows selective retention of diagnostically informative specimens, avoiding wasted effort on non-diagnostic individuals. Orientation classification enables specimens in the desired orientation to proceed immediately, while suboptimally oriented specimens can be repositioned for informative processing. Anatomical segmentation allows targeted processing of specimens displaying diagnostic characters or targeted analysis of specific anatomical regions in subsequent workflow steps. Comparative analysis of model architectures shows task-specific selection is crucial: a Convolutional Neural Network achieved an accuracy of 0.94 for orientation, a Vision Transformer achieved 0.88 for sex, and a U-Net precisely segmented nine anatomical regions with a mean IoU of 0.78. These results demonstrate that robotic imaging combined with deep learning provides a validated foundation for high-throughput, targeted specimen processing, maximising efficiency and utility for taxonomic and trait-based analyses, and supporting scalable, sustainable biodiversity workflows.

## 1. Introduction

The global decline in biodiversity has created an urgent need to accelerate the documentation of life on Earth, a task for which traditional methods are insufficient (Costello et al., 2013; Stork, 2018). Modern large-scale sampling techniques, such as Malaise traps for insects, generate enormous quantities of specimens, overwhelming the capacity of the few available taxonomic experts (Karlsson et al., 2020). This influx of material, often dominated by hyperdiverse and poorly understood groups, so-called "dark taxa" (Page 2016; Hartop et al. 2024, 2024), has created a severe "taxonomic impediment," where the rate of specimen collection far outpaces the rate of identification and description. The scale of this challenge has drawn attention beyond the scientific community: dark taxa are now being highlighted in public science communication as emblematic of the biodiversity crisis (e.g. Jones 2025). Their overwhelming dominance in global insect samples – just 20 families account for over half of the species and specimens in Malaise trap collections worldwide (Srivathsan et al., 2023) – underscores the urgency of developing scalable solutions to process and understand this hidden majority.

To overcome this challenge, the field is moving towards an integrated vision for biodiversity assessment that combines robotics, high-throughput sequencing (HTS), and artificial intelligence into a unified pipeline (Wägele et al., 2022). This emerging paradigm of "taxon-omics" envisions an automated workflow where specimens are first sorted by size (Ascenzi et al., 2025) and handled by robotic systems like the DiversityScanner (Wührl et al., 2022). These systems capture high-resolution images (Klug et al., 2024) that serve as the basis for AI-driven classification (Caruso et al., 2025; Shirali et al., 2024) and morphometric analysis (Shirali et al., 2025). This process complements modern molecular workflows, such as high-throughput DNA barcoding, where specimens are often sequenced first and morphologically validated later in a "reverse taxonomy" approach (Srivathsan et al. 2021; Hartop et al. 2022). The ultimate goal is to create a rich "digital voucher" for every specimen, where molecular data is linked to high-quality images and quantitative morphological data, providing a holistic record of biodiversity.

Robotics and high-throughput imaging alone are insufficient for efficient analysis; the value comes from processing specimens in a way that informs downstream steps. By extracting sex, orientation, and anatomical features from images, our workflow ensures that specimens can be efficiently prioritised for both processing and downstream analyses, maximising the utility of the collected data. The Phoridae (Diptera), a hyperdiverse family of small scuttle flies and a model dark taxon, highlight the scale of this challenge. Species identification in this group relies on taxonomic keys that require specimens to be viewed from standardized angles (e.g., dorsal, lateral), are often applicable to only one sex (typically males with diagnostic terminalia), and require specific characters to be visible. Manually sorting thousands of millimeter-long flies (or their images) by orientation, sex, and visible anatomical features is time-consuming and limits the scalability of automated workflows.

While recent computer vision studies have successfully automated single tasks like sex identification in medically important insects (Fraiwan et al., 2025; Kittichai et al., 2021; Tuda & Luna-Maldonado, 2020) or anatomical segmentation in model organisms (Le et al., 2020; Toulkeridou et al., 2023), these efforts have not yet been integrated. A significant gap remains in combining these components into a comprehensive pre-processing workflow suitable for the dark taxa that dominate biodiversity samples.

Here, we address this challenge by developing a foundational AI module to automate prioritisation for subsequent processing and analysis. Using Phoridae as a model system, our workflow performs three tasks on 2D specimen images:

1. **Orientation Classification:** Determines if a specimen is in a dorsal, ventral, left lateral, or right lateral view.
2. **Sex Classification:** Identifies the specimen as male, female, or undetermined.
3. **Body Part Segmentation:** Delineates nine key anatomical regions (e.g., head, thorax, terminalia).

These tasks enable specimens to be efficiently organised for downstream processing: specimens in the desired orientation and sex can proceed directly, while others can be repositioned or selectively retained for subsequent steps. Segmentation provides a foundation for future automated morphometrics, allowing the extraction of quantitative trait data directly from images. In this paper, we detail the workflow’s architecture, provide a comprehensive comparison of competing deep learning models, and validate its performance. We show that this automated approach is fast, accurate, and shows a capacity for handling ambiguous cases in a reliable manner, offering a powerful tool to accelerate the integration of morphology into next-generation biodiversity discovery.

## 2. Materials and Methods

### 2.1. Image Acquisition

The dataset consisted of 1,281 specimens of scuttle flies (Diptera: Phoridae) sourced from Malaise trap samples. Specimens were processed and imaged using two systems: the DiversityScanner, a robotic system for automated handling and imaging (Wührl et al., 2022), and the Entomoscope, an open-source photomicroscope (Wührl et al., 2024). Each specimen was individually imaged while submerged in ethanol. The system captured a focal stack of a minimum of five images at different depths, which were then fused into a single, high-contrast, all-in-focus composite image using Helicon Focus software (*Helicon Focus - Helicon Soft*, 2025).

A preprocessing step was applied to all images (Figure 1) to standardize the input for our models and remove uninformative background. A custom-trained object detection model, based on YOLOv8 architecture (Jocher et al., 2022/2023), was used to automatically identify the primary region of interest (ROI) containing the insect. The images were then cropped to this ROI, ensuring consistent framing and centering of the specimen for all subsequent tasks. The complete, annotated image dataset is publicly available at [Zenodo](https://zenodo.org/records/17252298).

**Figure 1.**
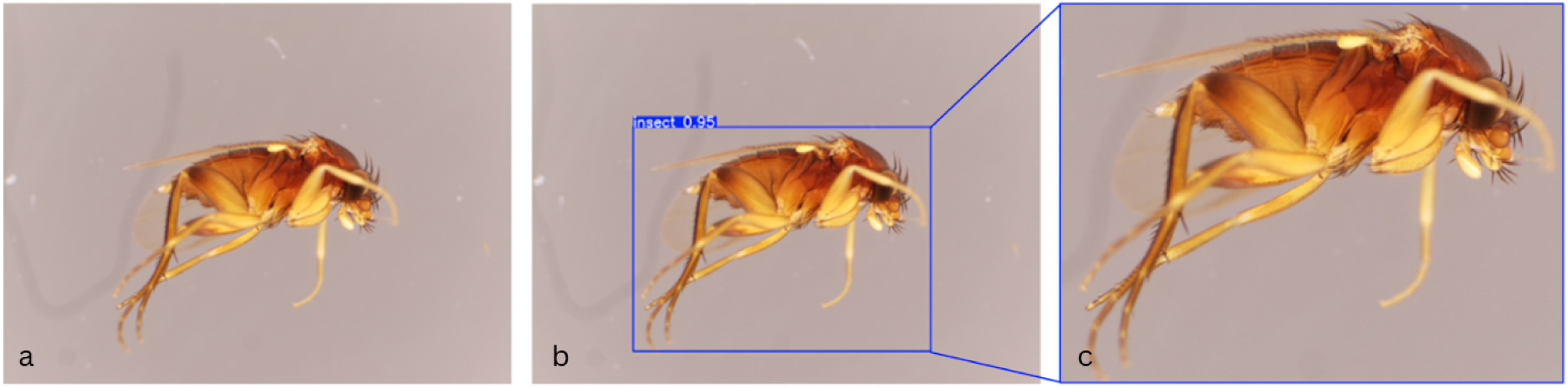
The automated image pre-processing pipeline. (a) A focus-stacked image of a Phoridae specimen. (b) A YOLOv8 model detects the insect and places a bounding box around the region of interest. (c) The final cropped image serves as the standardized input for all classification and segmentation models.

### 2.2. Ground-Truth Annotation

All images were annotated by taxonomic experts to create the ground-truth data for supervised learning. Annotations were performed using KAIDA, a specialized software tool for complex biological image annotation (Schilling et al., 2022). Three distinct annotation tasks were completed for each image:

- **Sex Classification:** Specimens were labeled as ‘Male’, ‘Female’, or ‘Undetermined’. The ‘Undetermined’ label was used when diagnostic features, primarily the terminalia (genitalia), were obscured by other body parts (e.g., legs, wings), out of focus, or otherwise not clearly visible to the expert.
- **Orientation Classification:** Specimens were assigned to one of four positional classes: ‘Dorsal’, ‘Ventral’, ‘Left Lateral’, or ‘Right Lateral’, based on the primary view presented in the image.
- **Body Part Segmentation:** Nine key anatomical regions were manually delineated with pixel-wise semantic masks: Head, Thorax, Abdomen, Antenna, Palps + labella, Legs, Wings, Scutellum, and Genitalia. These parts were chosen for their functional and diagnostic importance in Phoridae taxonomy.

The full dataset was split into training (70%), validation (15%), and test (15%) subsets. A stratified sampling approach was used to ensure that the proportional representation of each class within each task was maintained across the splits. This is particularly important for handling the inherent class imbalances in the dataset, such as the low number of specimens in a dorsal view. To prevent data leakage and ensure a fair evaluation, all images from a single specimen were kept within the same data split. The final distribution of images for each task is detailed in Table 1.

**Table 1.** Image count across tasks, classes, and data subsets (training, validation, and testing)

| Task | Class | Training | Validation | Test |
| --- | --- | --- | --- | --- |
| Sex Classification | Male | 590 | 126 | 127 |
|  | Female | 173 | 37 | 38 |
|  | Undetermined | 133 | 29 | 28 |
| Total (Sex Classification) |  | 896 | 192 | 193 |
| Orientation Classification | Dorsal | 47 | 10 | 10 |
|  | Left Lateral | 344 | 73 | 74 |
|  | Right Lateral | 310 | 67 | 66 |
|  | Ventral | 162 | 35 | 35 |
| Total (Orientation Classification) |  | 863 | 185 | 185 |
| Body Part Segmentation |  | 735 | 92 | 93 |

### 2.3. Deep Learning Architectures

To identify the optimal models for our workflow, we evaluated a range of deep learning architectures for the classification and segmentation tasks.

#### 2.3.1. Classification Models

For sex and orientation classification, we evaluated two major classes of deep learning models to compare their distinct architectural strategies. Convolutional Neural Networks (CNNs) (Lecun et al. 1998) are designed to detect local patterns (e.g., edges, textures) by applying filters that scan across an image. In contrast, Vision Transformers (ViTs) (Dosovitskiy et al. 2021) take a more holistic approach, treating an image as a sequence of patches and using attention mechanisms to model the relationships between them, allowing the capture of global context. Our comparison aimed to determine whether local feature extraction (CNNs) or global context modeling (ViTs) was more effective for our specific biological classification tasks. We tested architectures chosen to represent a broad spectrum of modern computer vision approaches:

- **Lightweight and Efficient CNNs:** We included YOLOv8 and YOLOv11 (Jocher et al., 2022/2023) as they are state-of-the-art models known for their balance of speed and accuracy, representing a practical choice for real-world deployment. For both architectures, we systematically trained all five standard model variants (from nano to x-large) and report the results for the best-performing ‘x-large’ variant for brevity.
- **High-Performance CNNs:** To explore the upper limits of CNN performance, we included ConvNeXt-XLarge (Liu et al., 2022) and EfficientNetV2-L (Tan & Le, 2019), which are larger, more complex architectures designed for maximum accuracy.
- **Vision Transformers (ViTs):** To test a fundamentally different approach, we included BEiTv2-large (Peng et al., 2022) and EVA-02-large (Fang et al., 2024). These models use global attention mechanisms instead of local convolutions, allowing them to capture broader contextual relationships within the image.

#### 2.3.2. Segmentation Model

For body part segmentation, we used the U-Net architecture, a standard for biomedical image segmentation due to its robust performance and efficient encoder-decoder structure with skip connections (Ronneberger et al., 2015). To optimize feature extraction within the U-Net, we experimented with two different backbones: EfficientNet-B0, known for its balance of efficiency and accuracy, and ResNet-18 (He et al., 2016), a well-established residual network.

### 2.4. Model Training and Implementation

All models were trained on the HAICORE high-performance computing cluster at the Karlsruhe Institute of Technology (KIT), using nodes equipped with NVIDIA A100 80GB GPUs. We used transfer learning for all models, initializing them with weights pre-trained on the ImageNet dataset (Deng et al., 2009) to accelerate training and improve generalization. The models were then fine-tuned on our Phoridae dataset using a comprehensive optimization strategy. Input image sizes were tailored to each model’s architecture, typically ranging from 224×224 to 640×640 pixels, to optimize performance. We applied a suite of data augmentation techniques to increase model robustness to variations in imaging conditions. These included photometric augmentations such as random changes in brightness, contrast, and saturation. For the segmentation and sex classification tasks, we also applied geometric augmentations, including horizontal flipping, scaling, and rotation. For the orientation classification task, geometric augmentations that would alter the orientation label (e.g., flipping and rotation) were explicitly excluded to avoid label ambiguity.

Our optimization strategy focused on achieving robust generalization and preventing overfitting. We used the AdamW optimizer for all classification models with an initial learning rate of 1e-4, and the Adam optimizer for segmentation with a learning rate of 1e-3. To address class imbalance, particularly for underrepresented classes like ‘Dorsal’ orientation, we applied class weights to the loss function for all classification tasks. For segmentation, we used a combined loss function of Focal Loss and Dice Loss, which is effective for handling extreme imbalances between foreground and background pixels. Furthermore, we implemented dropout regularization with a rate of 0.3 during training and employed an early stopping mechanism that halted training if validation loss did not improve for 15 consecutive epochs. Models were trained for a maximum of 150 epochs for classification and 250 epochs for segmentation. All models converged successfully (see Supplementary Figures S1 and S4 for training and validation curves for the best-performing models).

### 2.5. Evaluation

Model performance was assessed using a range of standard metrics on the held-out test set.

- **Classification:** We evaluated classification performance using Accuracy, Precision, Recall, and F1-Score. The F1-score, the harmonic mean of precision and recall, is particularly useful for datasets with class imbalance.
- **Segmentation:** Segmentation quality was measured using the Intersection over Union (IoU, or Jaccard index) and the Dice Score (an F1-score equivalent for segmentation). These metrics quantify the overlap between the predicted segmentation mask and the ground-truth annotation.
- **Independent Validation:** To benchmark our model against human-level performance, an independent expert taxonomist (not involved in the initial annotation) re-classified all images in the test set for both sex and orientation. This provided a direct comparison between model and expert accuracy. Additionally, to analyze the effect of feature visibility on segmentation performance, the independent tester annotated each target body part in the test images as ‘clearly visible’, ‘partially visible’, or ‘not visible’ based on its photographic clarity and focus. A detailed summary of the independent validation is provided in the Supplementary Material.

## 3. Results

### 3.1. Model Performance in Classification Tasks

Our comparative analysis of six architectures showed that the optimal model choice depended on the specific classification task (Table 2). For orientation classification, the CNN-based YOLOv8x-cls was the top-performing model, achieving an overall accuracy of 0.94 and a weighted F1-score of 0.89. In contrast, for sex classification, the Vision Transformer-based BEiTv2 model performed best, reaching an accuracy of 0.88 and the highest F1-score of 0.85. Notably, the ViT models (BEiTv2 and EVA-02) performed poorly on orientation classification (0.61 accuracy), suggesting their global attention mechanisms may be less effective than CNNs at distinguishing the local features that define left versus right lateral views.

**Table 2.** Model performance on classification tasks. Comparison of overall accuracy and weighted F1-score for six different model architectures on the test dataset. The best-performing model for each task is highlighted in **bold**. Scores are colour-coded to indicate performance levels: **green** (≥0.90), **yellow** (0.80–80.9), **orange** (0.70–0.79), and **red** (<0.70).

| Architecture | Sex Accuracy | Sex F1-Score | Orientation Accuracy | Orientation F1-Score |
| --- | --- | --- | --- | --- |
| YOLOv8x-cls | 0.88 | 0.82 | **0.94** | **0.89** |
| YOLO11x-cls | 0.82 | 0.75 | 0.90 | 0.86 |
| BEiTv2 | **0.88** | **0.85** | 0.61 | 0.72 |
| EVA-02 | 0.85 | 0.81 | 0.61 | 0.66 |
| ConvNeXtXLarge | 0.85 | 0.81 | 0.66 | 0.73 |
| EfficientNetV2L | 0.82 | 0.075 | 0.91 | 0.88 |

A detailed analysis of the best-performing models (Table 3) shows strong performance on the most common classes. A detailed breakdown of misclassifications for the top models is provided in the confusion matrices in the Supplementary Material (Figure S2). For orientation, the YOLOv8x-cls model achieved excellent precision and recall for Left Lateral (F1 = 0.96) and Right Lateral (F1 = 0.97) views. Performance was lower for the Ventral view (F1 = 0.90) and lowest for the Dorsal view (F1 = 0.74), corresponding to the small number of dorsal examples in the training set (n=56).

**Table 3.** Detailed classification report for the best-performing models. Per-class precision, recall, and F1-score for the YOLOv8x-cls model on the orientation task and the BEiTv2 model on the sex task. Scores are colour-coded to indicate performance levels: **green** (≥0.90), **yellow** (0.80–0.89), **orange** (0.70–0.79), and **red** (<0.70).

| Task | Class | Precision | Recall | F1-Score | Support |
| --- | --- | --- | --- | --- | --- |
| Orientation | Dorsal | 0.78 | 0.7 | 0.74 | 10 |
|  | Left Lateral | 0.95 | 0.97 | 0.96 | 74 |
|  | Right Lateral | 0.98 | 0.95 | 0.97 | 66 |
|  | Ventral | 0.89 | 0.91 | 0.9 | 35 |
|  | Accuracy |  |  | 0.94 | 185 |
|  | Macro Avg | 0.9 | 0.89 | 0.89 | 185 |
|  | Weighted Avg | 0.94 | 0.94 | 0.94 | 185 |
| Sex | Female | 0.94 | 0.87 | 0.9 | 38 |
|  | Male | 0.93 | 0.89 | 0.91 | 127 |
|  | Undetermined | 0.65 | 0.86 | 0.74 | 28 |
|  | Accuracy |  |  | 0.88 | 193 |
|  | Macro Avg | 0.84 | 0.87 | 0.85 | 193 |
|  | Weighted Avg | 0.89 | 0.88 | 0.88 | 193 |

For sex classification, the BEiTv2 model performed well on both Male (F1 = 0.91) and Female (F1 = 0.90) classes. The F1-score for the ‘Undetermined’ class was lower (0.74), which is expected given the inherent ambiguity of these images.

### 3.2. Model Performance in Body Part Segmentation

For the body part segmentation task, the U-Net architecture with an EfficientNetB0 backbone outperformed the ResNet18 backbone (Table 4). The model trained successfully, showing stable convergence of loss and IoU scores (Supplementary Figure S4), and achieved a mean IoU of 0.78 and a mean Dice score of 0.87 across all nine anatomical classes.

**Table 4.** Body part segmentation performance. Comparison of Intersection over Union (IoU) and Dice scores for the U-Net architecture with EfficientNetB0 and ResNet18 backbones on the test dataset. Scores are colour-coded to indicate performance levels: **green** (≥0.90), **yellow** (0.80–0.89), **orange** (0.70–0.79), and **red** (<0.70).

| Class | IoU<br>(EfficientNetB0) | Dice Score<br>(EfficientNetB0) | IoU (ResNet18) | Dice Score<br>(ResNet18) |
| --- | --- | --- | --- | --- |
| Head | 0.84 | 0.91 | 0.82 | 0.90 |
| Thorax | 0.84 | 0.91 | 0.83 | 0.91 |
| Abdomen | 0.85 | 0.92 | 0.83 | 0.91 |
| Antenna | 0.72 | 0.84 | 0.72 | 0.84 |
| Palps + labella | 0.71 | 0.83 | 0.72 | 0.84 |
| Legs | 0.82 | 0.90 | 0.81 | 0.89 |
| Wings | 0.88 | 0.93 | 0.87 | 0.93 |
| Halteres | 0.60 | 0.75 | 0.59 | 0.74 |
| Genitalia | 0.72 | 0.84 | 0.66 | 0.79 |
| Mean | 0.78 | 0.87 | 0.76 | 0.86 |

The model demonstrated excellent performance on large, well-defined body parts, with Dice scores above 0.91 for the Head, Thorax, Abdomen, and Wings. Performance was moderately lower for smaller or more complex structures like Legs (0.90), Antennae (0.84), and Genitalia (0.84). The lowest score was recorded for the Halteres (0.75), the smallest and most frequently obscured body part, which also had the lowest pixel representation in the dataset (see Supplementary Figure S3 for pixel count distribution).

### 3.3. Qualitative and Independent Validation

Visual inspection of the segmentation outputs aligns with the quantitative results, showing a high degree of accuracy and regional consistency (Figure 2). The example shown in Figure 2, which is representative of the model’s average performance, achieved an overall IoU of 0.79 and a Dice score of 0.87. The model’s predicted masks often appear smoother and more precise than the hand-labeled ground truth, particularly around complex boundaries. This suggests that the model learns generalized representations of anatomical structures, potentially surpassing the consistency of manual annotation.

**Figure 2.**
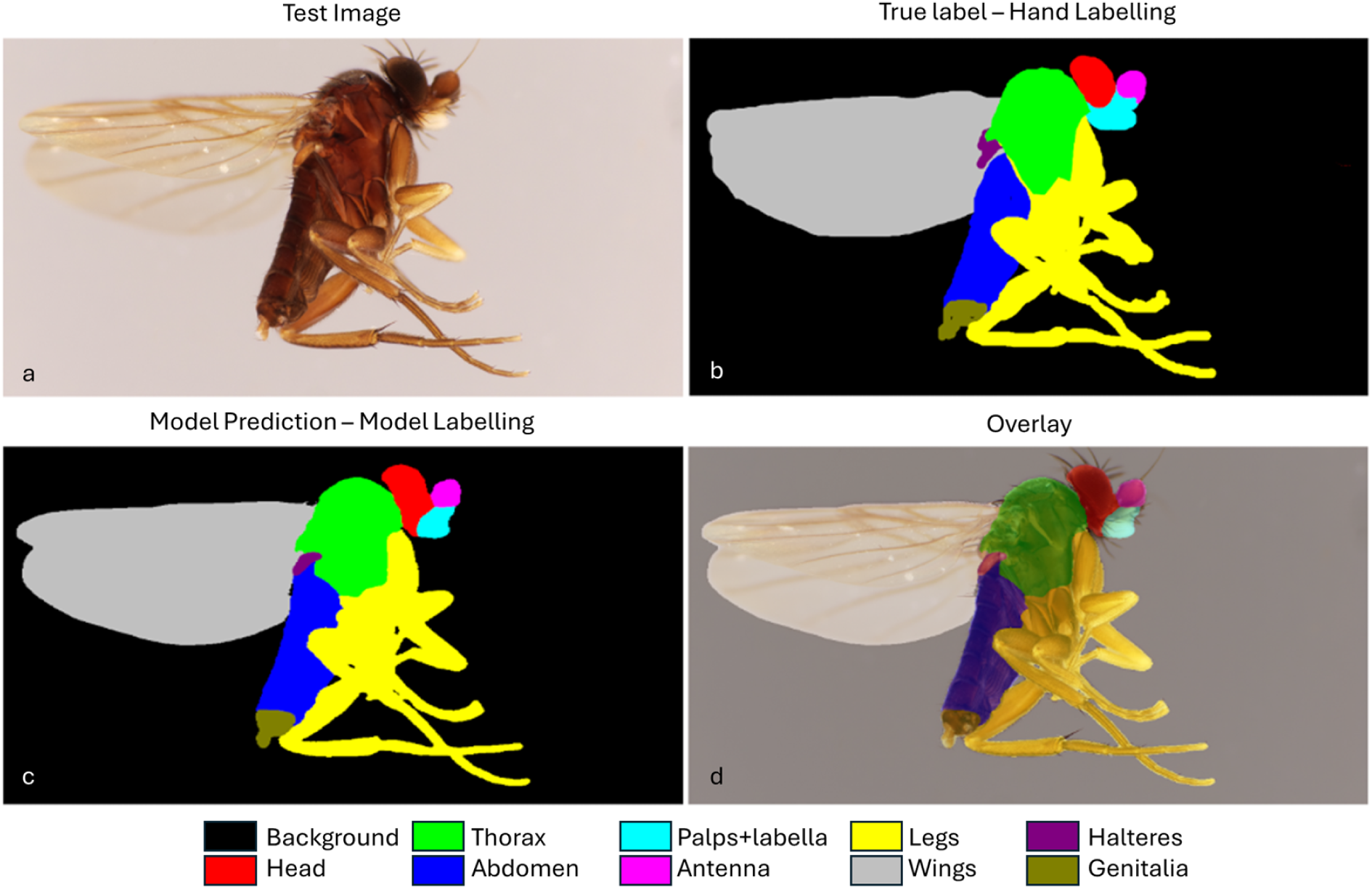
Qualitative results of body part segmentation. Comparison of (a) the original test image, (b) the hand-labeled ground-truth mask, (c) the model’s predicted segmentation mask, and (d) an overlay of the prediction on the original image. Each color corresponds to a different anatomical region, as shown in the legend. The prediction for this specific example achieved an overall IoU of 0.79 and a Dice score of 0.87, reflecting the model’s average performance.

The models also demonstrated a conservative but reliable strategy when dealing with challenging classification scenarios. The model’s performance on the ‘Undetermined’ class revealed a notable trend. While it assigned a sex to 4 of the 28 specimens that experts had labeled ‘Undetermined’ (Figure 3a), its more frequent error was to classify 13 known males as ‘Undetermined’, demonstrating a cautious behavior when faced with uncertainty. For specimens in ambiguous orientations (Figure 3b), the workflow can resolve these intermediate cases by providing both a primary and a valid secondary classification using a top-2 prediction strategy. Crucially, explainable AI techniques confirm that the models learn to focus on the correct anatomical regions; an Eigen-CAM visualization (Muhammad & Yeasin, 2020) shows the CNN-based orientation model focuses on the head and thorax (Figure 3c), while attention score maps from the Vision Transformer show the sex classification mode’s attention is concentrated on the terminalia when identifying a specimen (Figure 3d). The full results of the independent validation are summarized in Table 5, showing close agreement between the top models and expert performance.

**Figure 3.**
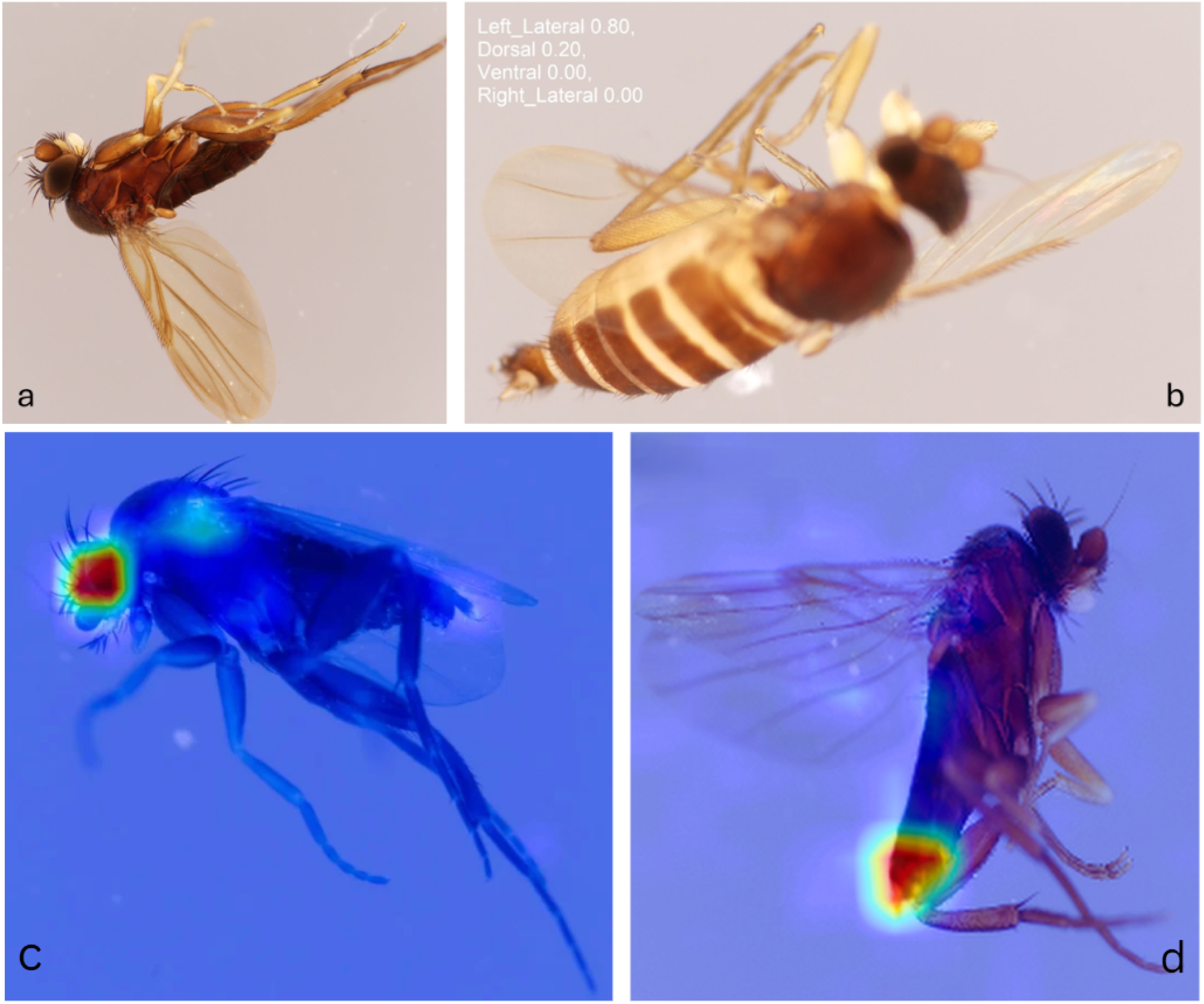
Analysis of challenging classification cases and model interpretability. (a) An example of the model assigning a sex (Male) to a specimen that experts had originally labeled as ‘Undetermined’. (b) A specimen in an ambiguous orientation between dorsal and lateral views, a case that can be handled by a logically constrained top-2 prediction strategy. (c) An Eigen-CAM visualization, showing the orientation model (red heatmap), is correctly focused on the head and thorax regions. (d) An attention score map from the Vision Transformer confirms that the sex classification model’s attention is concentrated on the terminalia when making predictions.

**Table 5.** Comparison of expert and model predictions. A summary of the agreement between the ground-truth labels, the top model predictions, and the independent expert’s classifications on the test set. Scores are colour-coded to indicate performance levels: **green** (≥0.90), **yellow** (0.80–0.89), **orange** (0.70–0.79), and **red** (<0.70).

| True Label | Total | Prediction<br>Matches | Prediction<br>Differences | Expert's<br>Matches | Expert's<br>Differences | Prediction<br>Accuracy | Expert's<br>Accuracy |
| --- | --- | --- | --- | --- | --- | --- | --- |
| Female | 38 | 33 | 5 | 38 | 0 | 0.87 | 1 |
| Male | 127 | 114 | 14 | 125 | 2 | 0.89 | 0.97 |
| Undetermined | 28 | 24 | 4 | 27 | 1 | 0.86 | 0.96 |
| Dorsal | 10 | 7 | 3 | 10 | 0 | 0.7 | 1 |
| Left Lateral | 74 | 72 | 2 | 74 | 0 | 0.972 | 1 |
| Right Lateral | 66 | 63 | 3 | 66 | 0 | 0.954 | 1 |
| Ventral | 35 | 32 | 3 | 35 | 0 | 0.914 | 1 |

## Discussion

We have developed and validated a multi-component deep learning workflow that extracts key biological information from specimen images, enabling targeted and efficient processing in the systematic pipeline. High accuracy across orientation classification, sex identification, and anatomical segmentation demonstrates that AI can reliably prioritise informative specimens and provide a robust foundation for downstream taxonomic and trait-based analyses. While adaptability to other taxa remains to be tested, this framework provides a foundation for developing more efficient and scalable taxonomic workflows across diverse organismal groups. Other researchers could apply this approach to their taxa by first generating high-quality specimen images (e.g., via robotic or semi-automated imaging), annotating a representative subset for key traits (sex, orientation, anatomical regions), and training task-specific deep learning models. Importantly, previous analyses of Malaise trap samples show that a small number of taxa dominate collections—approximately ten families account for roughly 50% of all specimens (Srivathsan et al., 2023). This suggests that a limited set of well-tuned models could capture a substantial fraction of collected material, providing an efficient route to scale up taxonomic processing across diverse taxa.

### 4.1. An Accurate and Automated Workflow for Early-Stage Specimen Sorting

This work introduces a deep learning workflow embedded in robotic specimen processing systems to automate three critical tasks—orientation classification, sex identification, and anatomical segmentation—from high-resolution insect images. By enabling targeted, high-throughput prioritisation of diagnostically informative specimens from hyperdiverse dark taxa, it lays the foundation for scalable, morphology-integrated biodiversity discovery. With accuracies of 0.94 for orientation and 0.88 for sex, our workflow is sufficiently accurate for practical implementation. In a high-throughput pipeline, this module could efficiently and automatically sort large samples, flagging just a small fraction for expert review. This represents a significant saving of expert time and a major step towards scaling up biodiversity research.

Our comparative analysis also provided a key insight into model selection: there is no one-size-fits-all architecture. The superiority of the CNN-based model for orientation classification suggests this task relies on identifying strong, local geometric features, a conclusion supported by XAI visualizations showing the model focusing on the head and thorax (Figure 3c). Conversely, the success of the ViT-based model on sex classification implies this task requires a more holistic assessment of the image, likely integrating subtle textural cues and global morphological patterns. The use of explainable AI (Figure 3d) provides strong evidence for this, confirming the model has learned to focus on the biologically relevant terminalia. This finding underscores the importance of empirical testing and task-specific model selection in applied AI for biology.

### 4.2. Beyond Sorting: The Foundational Role of Segmentation

The accurate segmentation of body parts is not an endpoint, but a gateway to a suite of downstream applications that are central to the future of digital systematics. The precise masks generated by our U-Net model can serve as the foundation for:

1. **Automated Morphometrics**: The pixel data from segmented regions like the head, thorax, and wings can be used to automatically calculate lengths, areas, and volumes. This enables the non-invasive extraction of key functional trait data at a scale impossible with manual methods.
2. **Automated Character Extraction:** The workflow could be extended to automatically score discrete morphological characters for phylogenetic matrices (e.g., presence/absence of certain bristles on the segmented thorax, or ratios of leg segments).
3. **Targeted Re-imaging:** In an integrated robotic pipeline, the segmentation output could direct the imaging system to perform a high-resolution scan of a specific region of interest, such as the terminalia, for detailed taxonomic study.
4. **Guiding AI-based Identification:** The segmentation output can act as a quality control filter for subsequent species identification models. For example, an identification model could be instructed not to process a specimen if the segmentation masks indicate that key diagnostic features, such as the terminalia, are not visible.

By providing this foundational data layer, our segmentation module is a critical technology for the next generation of quantitative and automated biodiversity analysis.

### 4.3. Model Robustness, Limitations, and Future Directions

Our models also demonstrated a conservative and reliable strategy when faced with ambiguous cases. An analysis of the sex classification model’s performance on the ‘Undetermined’ class provides a key insight. While the model successfully assigned a correct sex to 4 specimens that experts had found ambiguous (e.g., Figure 3a), its dominant behavior was to err on the side of caution. It misclassified 13 clear males as ‘Undetermined’, whereas it only made 6 direct misclassifications between the Male and Female classes combined. This tendency to classify uncertain cases as ‘Undetermined’ rather than making a definitive but potentially incorrect assignment is a valuable safety feature in an automated sorting pipeline, as it minimizes downstream errors by flagging difficult specimens for expert review. Furthermore, we developed a method to address the ambiguity of specimens positioned between two standard orientations. By implementing a logically constrained top-2 prediction system, the workflow can flag these intermediate cases for review. This system considers a secondary prediction only if it is biologically plausible (e.g., a ‘dorsal’ secondary prediction for a ‘lateral’ primary prediction) and its confidence exceeds a 5% threshold. This provides a robust solution to a common practical challenge without forcing a single, potentially incorrect, classification.

However, we acknowledge several limitations that suggest clear avenues for future work. Model performance was weakest on the most underrepresented classes, such as the ‘Dorsal’ orientation. For segmentation, the lowest scores were for ‘Halteres’, a classic challenge of data imbalance. This result is further explained by our independent visibility assessment, which revealed that halteres were labeled as ‘not visible’ or ‘partially visible’ in 34% of the test images (see Supplementary Table S1). This confirms that the model’s performance is logically constrained by the quality and visibility of features in the input data, not just by algorithmic limitations. Finally, this study was confined to a single insect family. While Phoridae represents a challenging test case, the generalizability of these models to other insect groups with different morphologies remains to be tested.

### 4.4. Broader Implications for the Future of Systematics

This work delivers a validated module for integrating deep learning into robotic biodiversity pipelines. Automating sex identification, orientation, and anatomical segmentation ensures that only specimens that are diagnostically informative are carried forward for downstream molecular or morphological analyses. These capabilities are crucial for tackling the vast numbers of undescribed “dark taxa” that dominate global sampling efforts. As such, workflows are adopted more broadly, and they promise to accelerate species discovery and contribute to the creation of comprehensive, high-quality digital biodiversity records.

## Conclusion

We present a deep learning workflow that automates three core processing tasks: orientation classification, sex identification, and body part segmentation. Validated on a taxonomically challenging group, this system achieves near-expert-level performance and is ready for integration into large-scale, automated biodiversity pipelines. By streamlining specimen processing triage, it marks a practical advance toward high-throughput, data-driven species discovery.

## Acknowledgments

Our work was supported by funding from the Center for Integrative Biodiversity Discovery at the Museum für Naturkunde Berlin and by grant #ZF4717901SK9 of the program Natural, Artificial and Cognitive Information Processing (NACIP) of the Helmholtz-Association, Germany; Helmholtz Association Initiative and Networking Fund on the HAICORE@KIT partition.

During the preparation of this work, the authors used AI tools to improve clarity and fluency in the writing process. After using these tools, the authors reviewed and edited the content as needed and take full responsibility for the content of the published article.

## Data Availability

All image data used in this study are available on [Zenodo](https://zenodo.org/records/17252298).

## Supporting Information

The full dataset from the independent expert validation, including the visibility assessment for each segmented body part, is available as a separate file: **Supplementary Data.**

### Supplementary Figures

**Figure S1.**
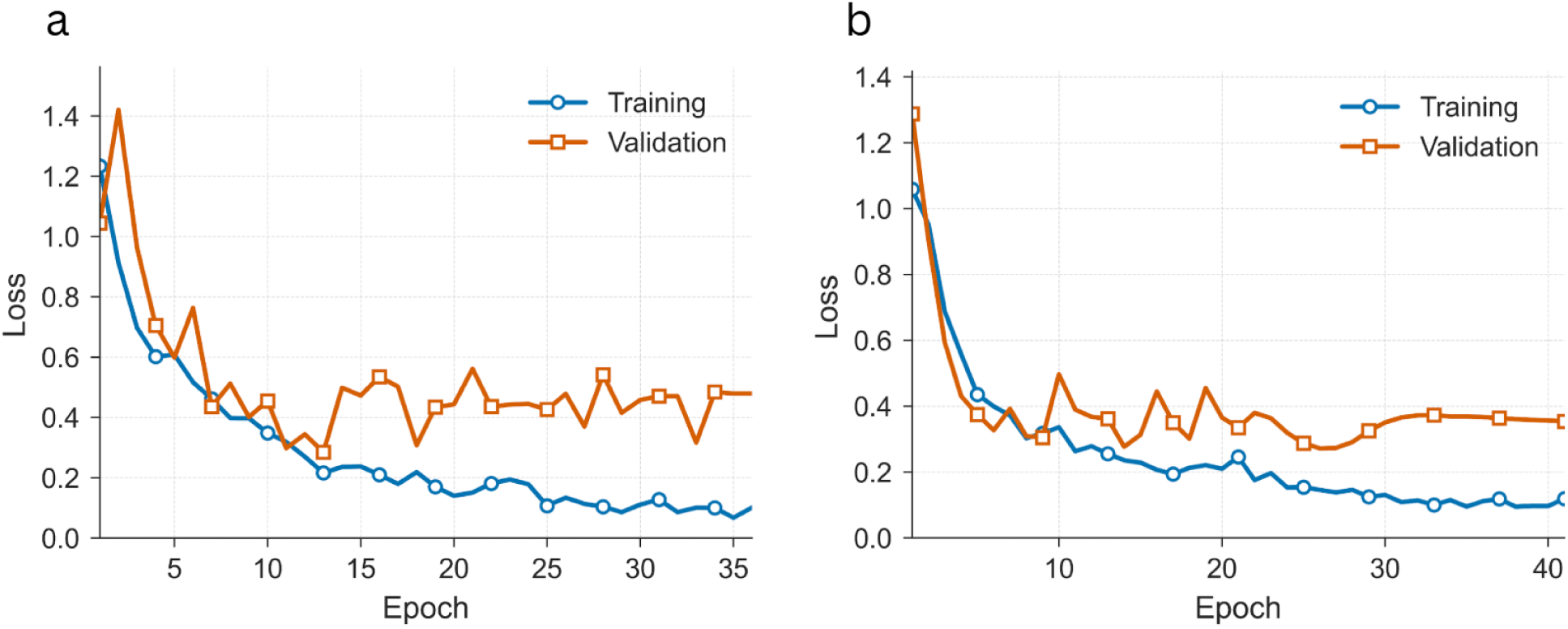
Training and validation curves for the best-performing models. (a) Loss curves for the YOLOv8x-cls model on the orientation classification task. (b) Loss curves for the BEiTv2 model on the sex classification task. The smooth convergence of the training and validation lines indicates that the models trained successfully without significant overfitting.

**Figure S2.**
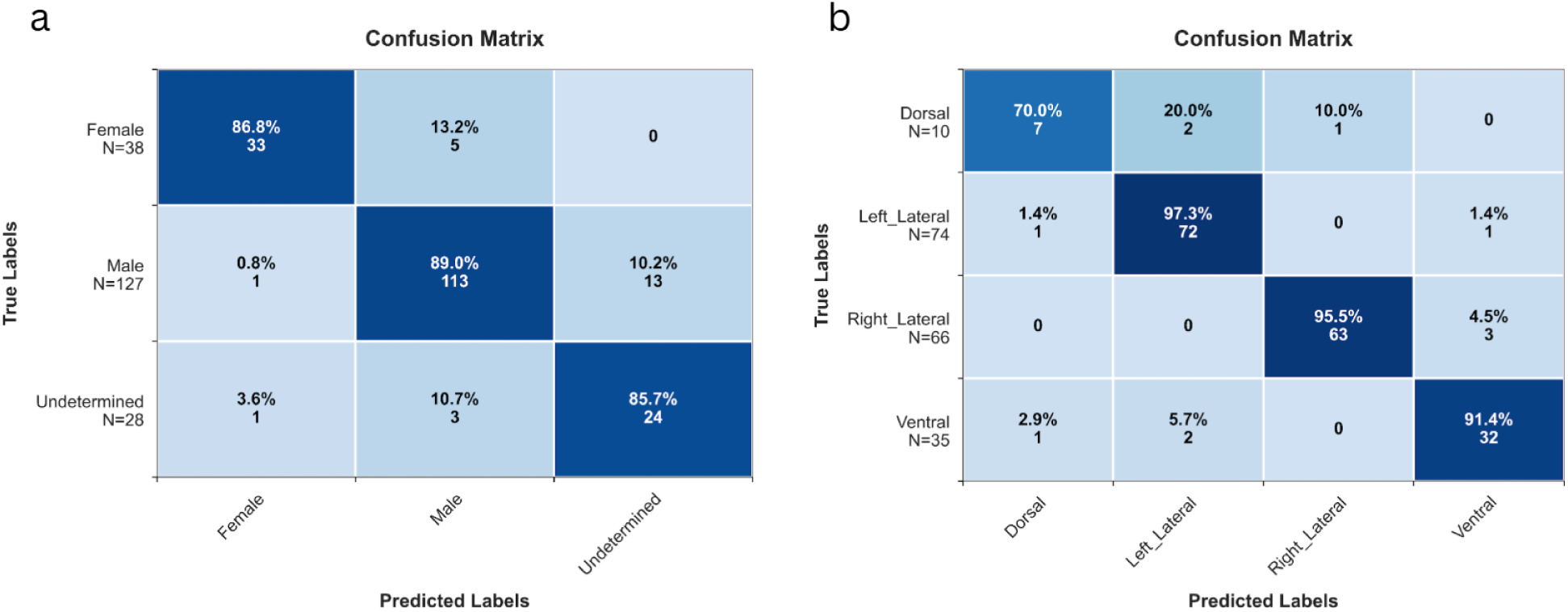
Confusion matrices for the best-performing classification models. (a) Confusion matrix for the BEiTv2 model on the sex task. (b) Confusion matrix for the YOLOv8x-cls model on the orientation task. Values on the diagonal represent correct classifications. Off-diagonal values show the number of specimens of a true class (y-axis) that were misclassified as a predicted class (x-axis).

**Figure S3.**
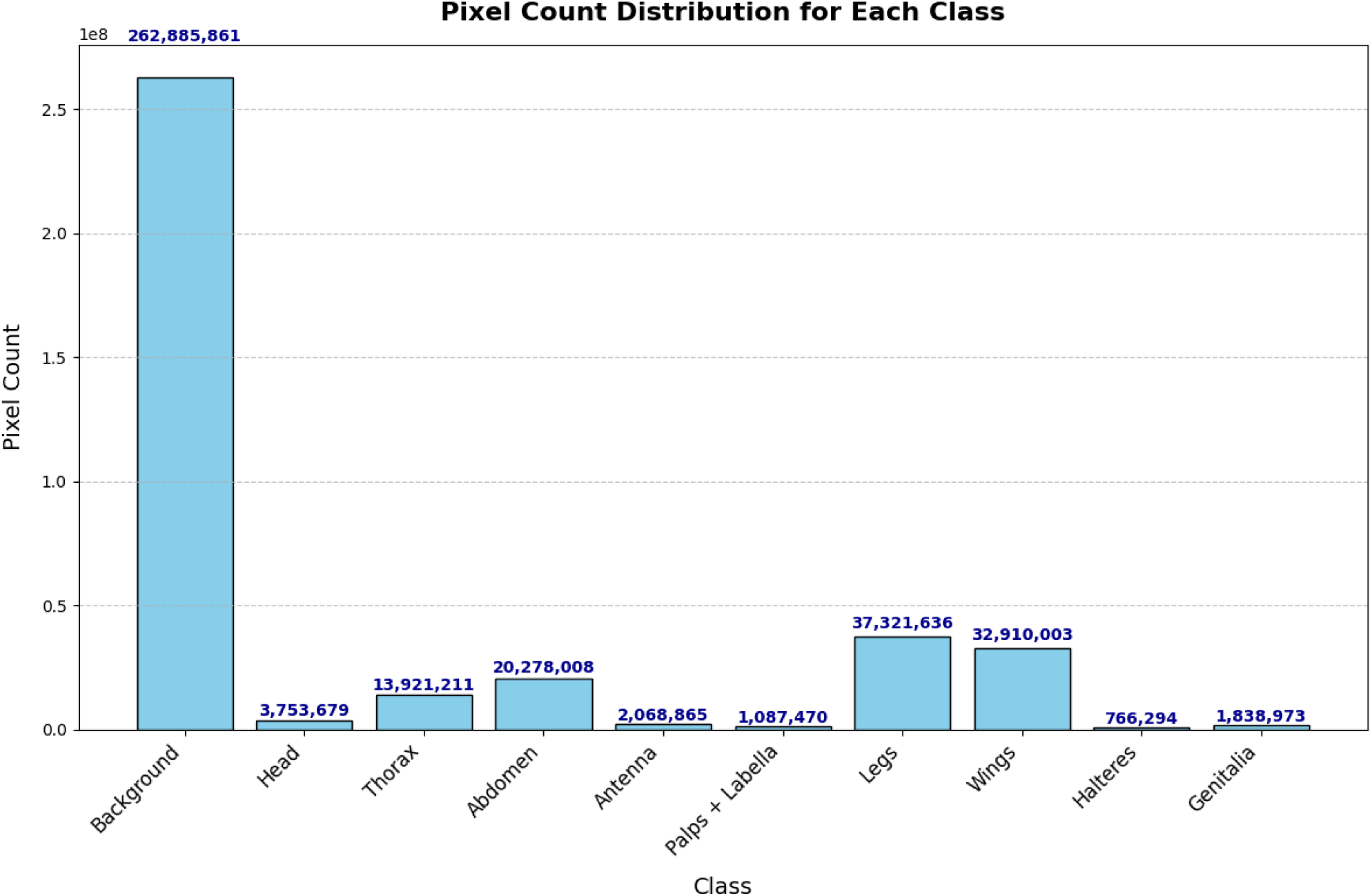
Pixel count distribution for each anatomical class in the segmentation dataset. The total number of pixels for each class across all training images highlights the significant class imbalance, with large regions like the wings and abdomen dominating the dataset, while small parts like the halteres are rare. This imbalance helps explain the lower segmentation performance on smaller body parts.

**Figure S4.**
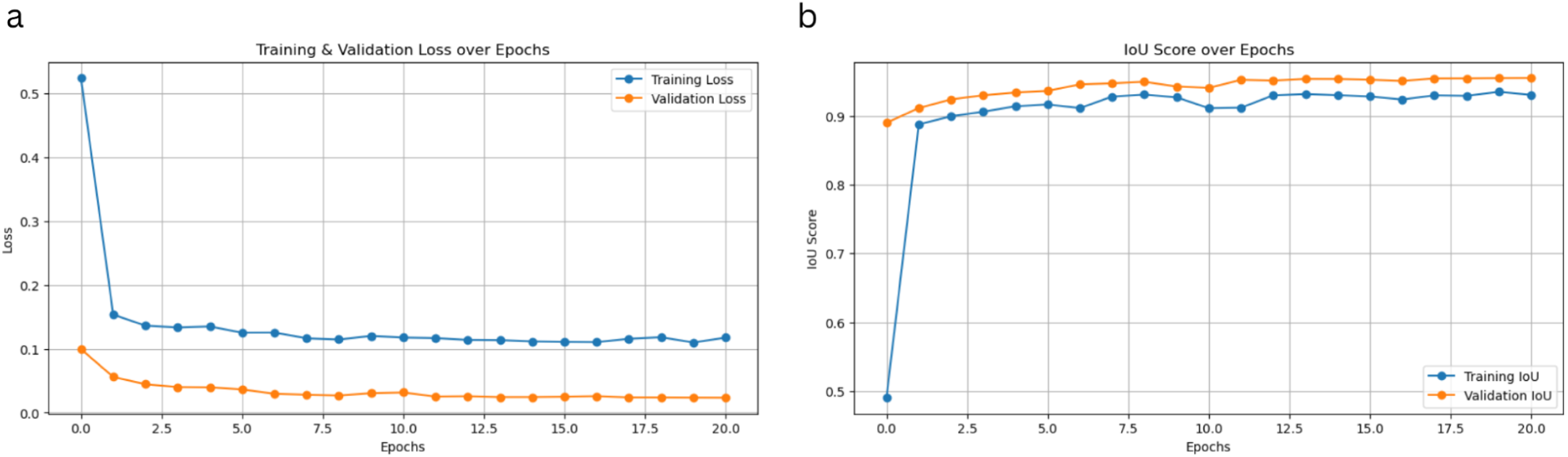
Training and validation curves for the best-performing segmentation model. (A) Loss and (B) Intersection over Union (IoU) curves for the U-Net with an EfficientNetB0 backbone over 21 training epochs. The smooth convergence of the training and validation lines indicates that the model trained successfully without significant overfitting.

